# Peripheral Beta-Blockade Differentially Enhances Cardiac and Respiratory Interoception

**DOI:** 10.1101/2025.02.28.640776

**Authors:** Ashley Tyrer, Jesper Fischer Ehmsen, Kelly Hoogervorst, Niia Nikolova, Victor Pando-Naude, Christian Holm Steenkjær, Arthur S. Courtin, Francesca Fardo, Tobias Hauser, Francesco Bavato, Micah Allen

**Affiliations:** Center of Functionally Integrative Neuroscience, Aarhus University, Denmark; Center for Music in the Brain, Aarhus University, Denmark; Department of Neurology, Aalborg University Hospital, Denmark; Institute of Neuroscience, Université catholique de Louvain, Belgium; Danish Pain Research Center, Aarhus University, Denmark; Department for Psychiatry and Psychotherapy, University of Tübingen, Germany; Max Planck UCL Centre for Computational Psychiatry and Ageing Research, University College London, United Kingdom; University Hospital of Psychiatry Zurich, University of Zurich, Switzerland; Cambridge Psychiatry, University of Cambridge, United Kingdom

**Keywords:** Interoception, Bayesian Modelling, Psychophysics, Respiratory, Cardiac, Noradrenaline, Metacognition

## Abstract

Interoception, the perception of internal visceral states, arises from complex brain-body interactions across the central and peripheral nervous systems. Despite noradrenaline’s key role in these interactions, its specific contribution to interoceptive processes remains unclear. In a placebo-controlled, randomised, within-subject study, we employed computational modelling of interoceptive psychophysics to determine how pharmacological beta-adrenoceptor antagonism controls interoception across cardiac and respiratory domains. Both cardio-selective bisoprolol and non-selective propranolol improved cardiac perceptual sensitivity, with bisoprolol exerting an enhanced effect on cardiac metacognition. In contrast, both beta-blockers increased respiratory perceptual precision, with no corresponding changes in sensitivity or metacognition. These findings reveal a novel dissociation between central and peripheral beta-adrenergic mechanisms in interoception, highlighting the pivotal role of peripheral noradrenaline in regulating multi-organ brain-body interactions. Our results suggest that beta-blockers may provide promising routes for modulating distinct facets of interoception, potentially opening new avenues for intervention in conditions characterised by disrupted bodily self-awareness.

## Introduction

Interoception is the ability to perceive and process signals related to the body’s internal state, such as those arising from the heart, lungs, and other visceral organs^1,2^. This processing is fundamental to emotional regulation and self-awareness, and its dysfunction is widely implicated in psychiatric conditions such as anxiety, depression, and panic disorder^3,4^. Despite this clear importance, the neurobiological - and particularly the neuropharmacological - mechanisms underlying interoception remain poorly understood. Disentangling these mechanisms is a critical step toward understanding how interoceptive disturbances contribute to mental health conditions and how they may be targeted for intervention.

Noradrenaline is a key neuromodulator in both the central and peripheral nervous systems that is well-positioned to modulate interoceptive processing. Centrally, the locus coeruleus regulates noradrenaline release, which controls neural computations governing perceptual and cognitive uncertainty^5–7^. This link between noradrenaline and uncertainty is an ideal target for pharmacological manipulation. For example, the centrally-acting beta-blocker propranolol has been found to modulate perceptual decision-making, metacognition, and learning^8–10^. In the peripheral nervous system, postganglionic sympathetic nerve terminals and the adrenal medulla release noradrenaline, where it binds to beta-adrenoceptors in the heart, lungs, and other visceral organs^11^. This signalling regulates heart rate, blood pressure, and respiratory function, modulating vagal tone and sympathovagal balance as part of the autonomic fight-or-flight response^12,13^. Given its dual role in central and peripheral autonomic regulation, noradrenaline may influence interoception through both cortical and visceral pathways, potentially controlling multiple dimensions of bodily signal processing^14–18^.

While much of interoception research has focused on cardiac signals, recent findings highlight respiratory interoception as a physiologically distinct yet understudied domain^19–21^. Noradrenaline is likely to influence interoception through organ-specific pathways, as β₁-adrenoceptors are predominantly concentrated in the heart, whereas β₂-adrenoceptors play a greater role in pulmonary function^22,23^. By exploiting the differential affinities of selective versus non-selective beta-blockers, we can determine whether beta-adrenergic pathways exert general effects across interoceptive domains or instead differentially modulate interoceptive processes according to the organ system they most directly regulate. Understanding these differences may have unique implications for how we understand psychological disorders, as cardiac and respiratory interoception are also uniquely associated with various symptom profiles^3,24–29^.

To determine whether peripheral and central beta-adrenergic pathways differentially influence interoception across cardiac and respiratory domains, we compared propranolol, a non-selective beta-blocker, with bisoprolol, a highly cardioselective beta-blocker. We hypothesised that if central beta-adrenergic pathways are pivotal for interoception, propranolol would exert the strongest effects regardless of domain. Conversely, if peripheral signalling alone is sufficient, bisoprolol should produce an equivalent or greater effect. To test these hypotheses, we utilised hierarchical Bayesian modelling of interoceptive psychophysics to assess whether these beta-blockers selectively modulate interoceptive sensitivity, precision, or metacognition across physiological domains. Both bisoprolol and propranolol enhanced cardiac perceptual sensitivity, with bisoprolol exerting a stronger effect on cardiac metacognition. In contrast, both beta-blockers increased respiratory perceptual precision, without altering sensitivity or metacognition. These findings provide novel insights into how distinct neuromodulatory pathways regulate brain-body interactions, highlighting the potential to selectively target interoceptive processes through pharmacological intervention.

## Results

To differentiate the roles of central and peripheral noradrenaline in interoception, we conducted a randomised, double-blind, placebo-controlled, within-subject study in 50 healthy young adults (see **Figure 1A** for study outline). Participants received either propranolol, a non-selective beta-blocker that antagonises both β_1_- and β_2_-adrenoceptors and readily crosses the blood-brain barrier, or bisoprolol, a highly β_1_-selective antagonist with minimal central infiltration^30,31^. Given propranolol’s broad adrenergic inhibition and central effects, it allowed us to assess the contribution of both peripheral and central pathways, whereas bisoprolol’s cardioselectivity and limited β_2_ binding enabled us to isolate the effects of targeted peripheral beta-adrenergic modulation. Consequently, we determined whether interoceptive processing is modulated by peripheral signalling alone or requires central noradrenergic involvement.

**Figure 1:**
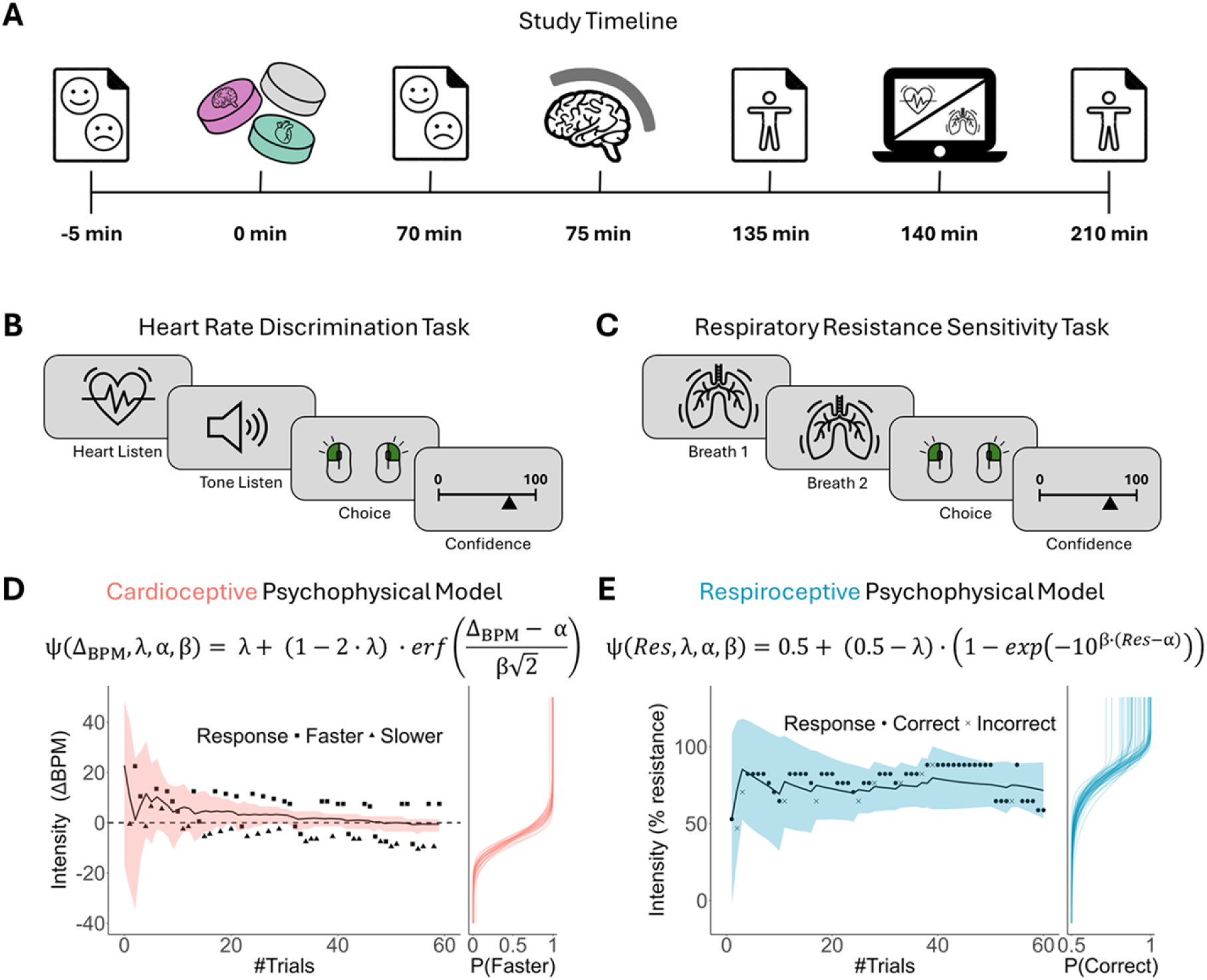
Study Design and Multimodal Interoceptive Sensitivity Battery. (**A**) Study visit timeline displaying baseline measures, drug administration, and interoceptive tasks. Participants were administered a single oral acute dose of either 40 mg propranolol, 10 mg bisoprolol, or a matched placebo. Immediately before and 70 mins following drug administration, participants completed the Positive And Negative Affect Scale (PANAS) and STICSA self-report surveys. Participants also completed the Interoceptive Belief Questionnaire (IBQ) before and after the interoceptive tasks. (**B**) Heart Rate Discrimination Task (HRDT). Participants compared auditory tones to their real-time heart rate, monitored using a pulse oximeter, and provided confidence ratings following their decisions. (**C**) Respiratory Resistance Sensitivity Task (RRST). Participants inhaled twice through a respiratory circuit, judging which of the two breaths they found to be more difficult and provided confidence ratings following their decisions. (**D**) Psychophysical model for the HRDT. *Top:* equation for cardioceptive psychophysical model, estimating sensitivity (threshold, α), precision (slope, β), and lapse rate (λ). *Left:* adaptive adjustments of tone frequency (ΔBPM) across 60 trials in an example participant. *Right:* example psychometric fit. (**E**) Psychophysical model for the RRST. *Top:* equation for respiroceptive psychophysical model, estimating sensitivity (threshold, α), precision (slope, β), and lapse rate (λ). *Left:* adaptive adjustments of respiratory resistance (% resistance) across 60 trials in an example participant. *Right:* example psychometric fit. Example data demonstrate the adaptive stimulus adjustments of both tasks on a single-subject level.

To independently assess cardiac and respiratory interoception, we employed two validated psychophysical tasks (**Figure 1B-C**): the Heart Rate Discrimination Task (HRDT) and the Respiratory Resistance Sensitivity Task (RRST)^32,33^. In the HRDT, participants classified an auditory tone as either “faster” or “slower” than their current heart rate. The RRST similarly required participants to compare two successive breaths and identify which contained a resistive load delivered via a computer-controlled inspiratory circuit. For both tasks, the stimuli (i.e., the tempo of auditory tones in the HRDT, and magnitude of resistive load in the RRST) were dynamically adjusted using a Bayesian adaptive algorithm^34^.

We then applied hierarchical Bayesian modelling to estimate the following psychometric parameters: threshold (interoceptive sensitivity, α), slope (interoceptive precision, β), and lapse rate (λ) (**Figure 1D-E**). The threshold represents the smallest detectable change in an interoceptive signal, with lower values indicating greater sensitivity to subtle internal fluctuations. A steeper slope, in turn, indicates greater precision, as participants more reliably differentiate near-threshold stimuli with minimal uncertainty. Lapse rate then captures the proportion of random or inattentive responses, allowing us to distinguish perceptual performance from errors due to lapses in attention or motor execution. We further quantified metacognitive awareness as the alignment between trial-wise confidence and choice accuracy^35–37^. These methodological advances enabled us to derive robust, interpretable estimates of interoceptive sensitivity, precision, and metacognition across physiological domains^38,39^.

### Reduced mean heart rate and increased heart rate variability under beta-blockade

As a positive control analysis to verify the cardiac effects of each drug manipulation, we first assessed the effects of propranolol, bisoprolol, and placebo on mean heart rate and heart rate variability (HRV) through analysis of resting-state electrocardiogram (ECG) data (mean heart rate (BPM ± SD): placebo = 65.7 ± 9.67; propranolol = 55.7 ± 8.19; bisoprolol = 55.0 ± 8.35). In line with the above analysis, mean heart rate was significantly reduced under both propranolol and bisoprolol compared to placebo (propranolol: *t*(46) = 11.5, *p* < .001; bisoprolol: *t*(45) = 10.8, *p* < .001). There was no significant difference in mean heart rate between propranolol and bisoprolol (*t*(46) = 1.59, *p* = .118) (**Supplementary Figure S1A**). HRV (RMSSD) was also significantly increased under propranolol and bisoprolol compared with placebo (propranolol: *t*(46) = −5.92, *p* < .001; bisoprolol: *t*(45) = −6.15, *p* < .001), and HRV was also significantly increased under bisoprolol compared with propranolol (*t*(46) = −2.25, *p* = .029) (**Supplementary Figure S1B**). Finally, sympathovagal balance (low frequency to high frequency (LF/HF) ratio) was significantly reduced under propranolol and bisoprolol compared with placebo (propranolol: *t*(46) = 4.33, *p* < .001; bisoprolol: *t*(45) = 4.21, *p* < .001). There was no significant difference in LF/HF ratio between propranolol and bisoprolol (*t*(46) = 0.032, *p* = .975) (**Supplementary Figure S1C**). These results indicate that both propranolol and bisoprolol robustly reduce sympathetic tone while enhancing parasympathetic influence, as evidenced by reduced heart rate and elevated HRV. The heightened HRV under bisoprolol in particular aligns with its more selective peripheral action, pointing to a distinct physiological profile of peripheral beta-adrenoceptor blockade.

### No effect of beta-blockade on mood, affect, or somatic anxiety

To determine whether potential interoceptive effects were driven by shifts in mood or anxiety, we assessed self-reported affect and somatic anxiety at baseline and 70 mins post-drug administration. Our analysis found no significant differences between drug conditions at either time point; affect and somatic anxiety scores all remained stable following drug administration (repeated-measures ANOVA; all contrasts: *p* > .05). These findings indicate that neither propranolol nor bisoprolol elicited measurable changes in acute affective states, and that any interoceptive modulations arise independently of changes in mood or somatic anxiety. Additionally, we examined the role of subjective beliefs in interoception using the Interoceptive Belief Questionnaire (IBQ), finding no meaningful effect of either drug on interoceptive beliefs or affect (see **Supplementary Methods S1** for analysis, **Supplementary Table 1** for IBQ items).

### Significant increase in cardioceptive threshold under both propranolol and bisoprolol

We investigated the effects of propranolol and bisoprolol in the HRDT and RRST using hierarchical Bayesian psychometric models, each tailored to the respective adaptive task (see Methods and Figure 1D-E). Estimating effects fully hierarchically allowed us to leverage partial pooling, improving statistical efficiency by reducing noise in individual estimates while preserving meaningful subject-level variation. This approach also accounted for random effects and baseline differences, ensuring that drug effects were estimated with greater precision while controlling for within-subject variability. In addition, all hierarchical HRDT models controlled for the effect of beta blockade on heart rate (see **Supplementary Methods S2-3**). Model fits were validated at the individual level (**Supplementary Figures 2–3**), confirming their robustness and reliability.

In the HRDT, we observed a significant increase in interoceptive threshold (α) under both propranolol (µ = 2.82, CI [0.57; 5.03], *P*(placebo > propranolol) = .02) and bisoprolol (µ = 2.51, CI [0.11; 4.88], *P*(placebo > bisoprolol) = .04), indicating improved sensitivity to heart rate under beta-blockade (**Figure 2A, C**). In contrast, no significant effects were observed on the slope (β) for propranolol (µ = −0.007, CI [−0.23; 0.20], *P*(placebo > bisoprolol) = .50) or bisoprolol (µ = −0.03, CI [−0.17; 0.12], *P*(placebo > bisoprolol) = .61), suggesting that beta-blockade did not alter cardioceptive precision. Similarly, no significant differences in lapse rate (λ) were found between propranolol (µ = 0.48, CI [−2.34; 2.94], *P*(placebo > bisoprolol) = .34) or bisoprolol (µ = −1.19, CI [−3.88; 1.34], *P*(placebo > bisoprolol) = .77) compared to placebo, indicating no change in the rate of random responses or button press errors under the active drug conditions. These results indicate that beta-blockade selectively enhances cardiac interoceptive sensitivity, as reflected by a shift in the discrimination threshold towards zero, without affecting the consistency of responses or error rates. In other words, propranolol and bisoprolol improve the ability to detect subtle changes in heart rate, while leaving precision and response biases intact.

**Figure 2.**
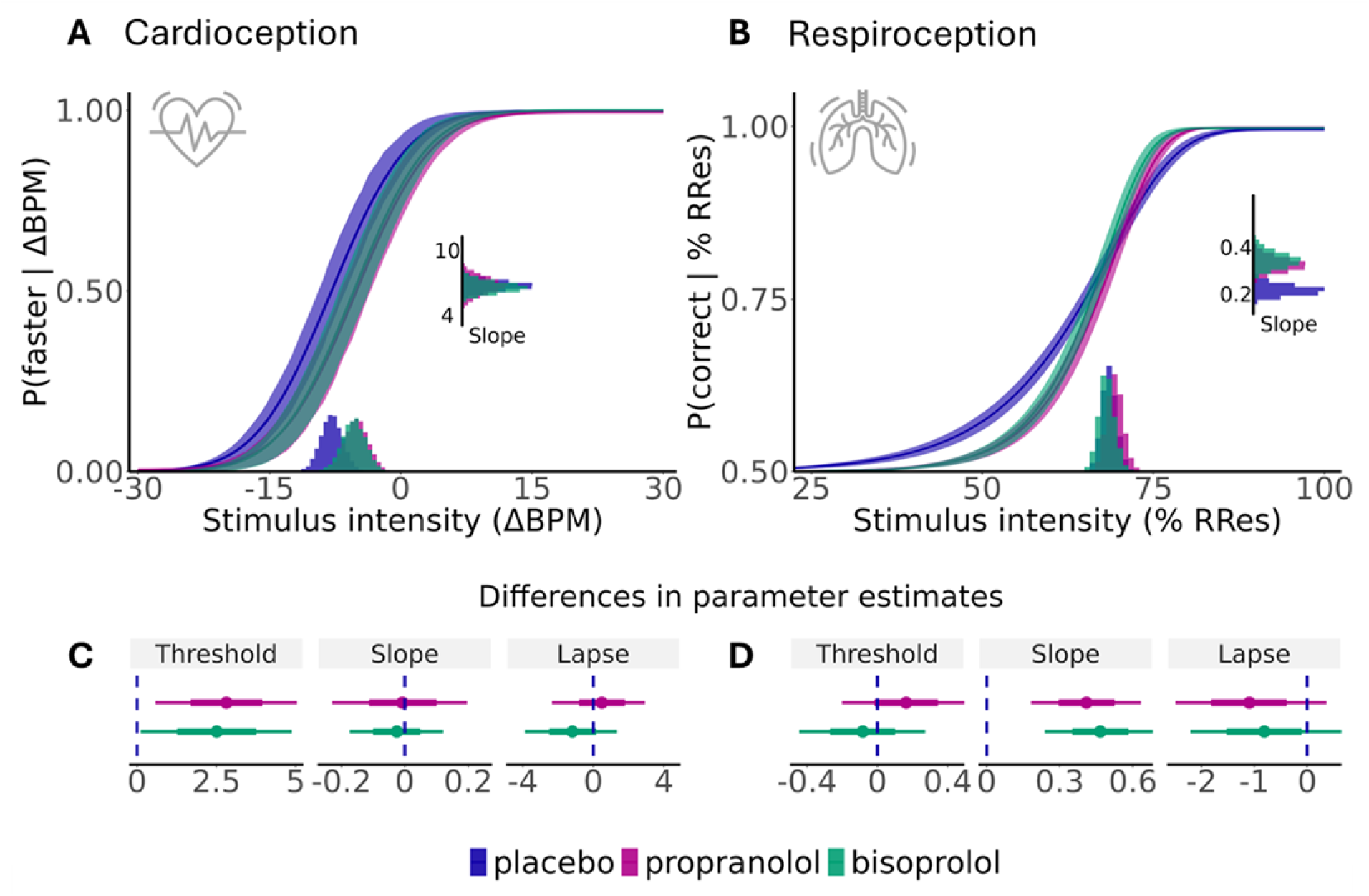
Psychophysical modelling of beta-blockade on cardioceptive and respiroceptive sensitivity (threshold, ɑ), and precision (slope, β). (**A**) **Beta blockade increased cardioceptive sensitivity.** Posterior predictive psychometric functions for each drug condition (placebo, propranolol, and bisoprolol) as a function of stimulus intensity (ΔBPM). Shaded regions represent 80% credible intervals (CI), highlighting differences in cardioceptive sensitivity across drug conditions. *Bottom*: posterior distribution of the difference in threshold values for all drug conditions, indicating a significant effect on cardioceptive thresholds for both contrasts (*P*(placebo > propranolol) = .02; *P*(placebo > bisoprolol) = .04). *Right:* Posterior distributions of slope estimates for all drug conditions, revealing no significant effects on cardioceptive precision. (**B**) **Beta blockade increases respiroceptive precision.** Posterior predictive psychometric functions for each drug condition as a function of stimulus intensity (% RRes), with 80% credible intervals shown. *Bottom:* posterior distributions of threshold estimates for placebo, propranolol, and bisoprolol, revealing no significant effects on respiroceptive thresholds. *Right:* posterior distributions of slope estimates for all drug conditions, revealing significantly increased respiroceptive precision for both drug conditions compared to placebo (*P*(placebo > propranolol) = .002; *P*(placebo > bisoprolol) = .0004). (**C**) **Cardioceptive parameter differences.** Posterior distributions of differences between placebo and drug conditions displayed with 80 and 95% CI for all three psychometric parameters for the HRDT. (**D**) **Respiroceptive parameter differences.** Posterior distributions of differences between placebo and drug conditions displayed with 80 and 95% CI for all three psychometric parameters for the RRST.

### Greater Respiroceptive Perceptual Precision under Central and Peripheral Beta-Blockade

We next applied our modelling approach to the RRST to assess the effects of beta-blockade on respiratory interoception. Our analysis revealed that slope estimates were significantly higher under both propranolol (µ = 0.41, CI [0.18; 0.63], *P*(placebo > propranolol) = .002) and bisoprolol (µ = 0.47, CI [0.24; 0.69], *P*(placebo > bisoprolol) = .0004) compared to placebo, indicating improved respiroceptive precision (**Figure 2B, D**). No significant differences were found for threshold (µ = 0.16, CI [−0.20; 0.52], *P*(placebo > propranolol) = .23; = −0.08, CI [−0.44; 0.27], *P*(placebo > bisoprolol) = .65) or lapse rate (propranolol: µ = −1.23, CI [−2.49; 0.39], *P*(placebo > propranolol) = .90; bisoprolol: µ = −0.80, CI [−2.20; 0.72], *P*(placebo > bisoprolol) = .83), suggesting that beta-blockade did not affect respiroceptive sensitivity or random response tendencies. Overall, these results indicate that beta-blockade selectively enhances the precision of respiratory signal processing, while leaving detection thresholds and response consistency intact.

### Improved Cardiac Metacognition under Central and Peripheral Beta-Blockade

To quantify metacognitive awareness, we fitted a hierarchical ordered beta regression model to examine how propranolol and bisoprolol influence the alignment of interoceptive confidence with choice accuracy, while controlling for mean resting heart rate. Heart rate exerted a significant positive effect on confidence (β = 1.22, CI [1.14; 1.31], *p* < .001), indicating that participants were more confident as their actual heart rate increased. As expected, discrimination accuracy was also positively associated with confidence (β = 0.69, CI [0.59; 0.79], *p* < .001). We found that both drugs increased metacognitive awareness, as indicated by a significant drug-by-response accuracy interactions, with propranolol (propranolol × accuracy: β = 0.88, CI [0.79; 1.00], *p* = .045) and bisoprolol (bisoprolol × accuracy: β = 0.85, CI [0.75; 0.96], *p* = .008) influencing confidence for correct versus incorrect responses (**Figure 3A**). Post-hoc tests revealed no significant differences between drugs for incorrect responses (all *p* > .05). However, both propranolol (z = −3.43, *p* = .002) and bisoprolol (z = −6.0, *p* < .001) significantly increased confidence compared to placebo for correct responses, with bisoprolol exerting the strongest effect (propranolol vs. bisoprolol: z = −3.23, *p* = .0035) (**Figure 3C**). These results indicate that beta-blockade enhances metacognitive awareness by improving the ability to accurately assess interoceptive performance, particularly by elevating confidence for correct choices.

**Figure 3.**
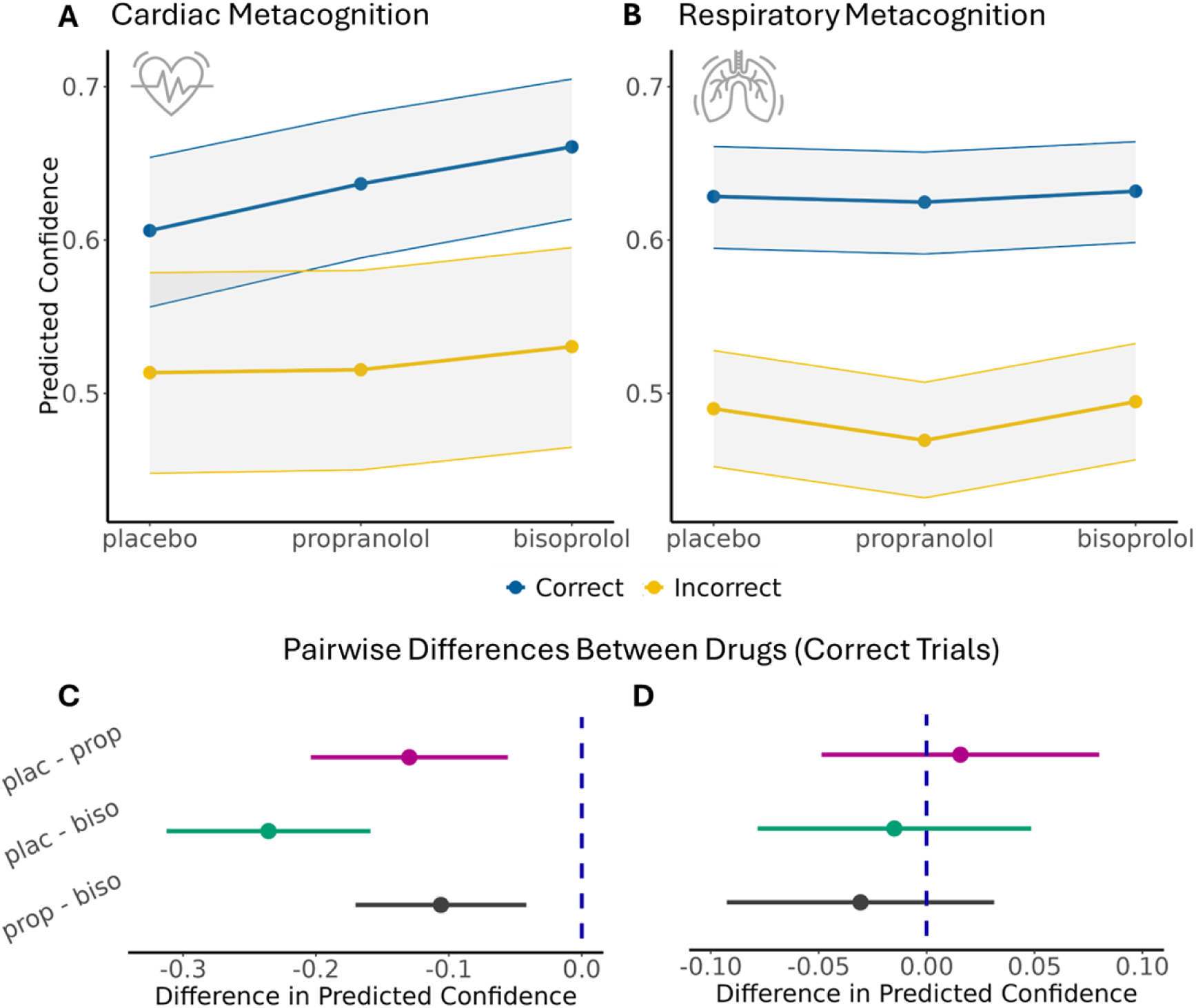
Ordered Beta Regression of Metacognition in HRDT Task (left) and RRST (right). Predicted confidence by drug and response accuracy in the HRDT (**A**) and RRST (**B**). Shaded area = 95% confidence intervals (CIs). Pairwise differences in predicted confidence between drugs during correct trials in the HRDT (**C**) and the RRST (**D**). Error bars = 95% CIs.

### No effect of beta-blockade on respiroceptive metacognition

To probe whether beta-blockade influences metacognitive awareness during respiratory interoception, we employed a separate hierarchical ordered beta regression while controlling for stimulus intensity (i.e., scaled respiratory resistance at each trial). As expected, stimulus intensity exerted a significant positive effect on confidence (β = 0.28, CI [0.24; 0.31], *p* < .001), with participants reporting greater confidence as the stimulus resistance increased. Accuracy was also strongly associated with higher confidence (β = 0.57, CI [0.45; 0.68], *p* < .001), reflecting robust metacognitive awareness. Neither propranolol (β = −0.07, CI [−0.19; 0.05], *p* = .26) nor bisoprolol (β = 0.17, CI [−0.1; 0.14], *p* = .77) exhibited significant main effects on confidence, and there were no significant interactions between drug condition and accuracy (propranolol × accuracy: β = 0.05, CI [−0.08; 0.19], *p* = .42; bisoprolol × accuracy: β = 0.008, CI [−0.13; 0.14], *p* = .90). These results indicate that, although stimulus resistance and choice accuracy robustly drive confidence judgments, beta-adrenoceptor antagonism does not modulate the metacognitive awareness of respiratory interoception (**Figure 3B, D**).

## Discussion

In this study, we demonstrate that central and peripheral noradrenergic beta-blockade exerts distinct effects on interoceptive sensitivity, precision, and metacognition across cardiac and respiratory domains. Both beta-blockers enhanced cardiac interoception, with bisoprolol showing a stronger effect on metacognitive awareness, while respiratory interoception was modulated through increased precision without changes in sensitivity or metacognition. These results reveal a domain-specific dissociation in the mechanisms by which noradrenaline influences interoception, and highlights distinct autonomic pathways that may be potential targets of future therapeutic interventions for psychiatric disorders.

For both the cardiac and respiratory interoceptive tasks, participants exhibited significantly altered interoceptive threshold (HRDT) and slope (RRST) under both propranolol and bisoprolol compared with placebo, but these parameters did not differ significantly between the two beta-blockade pathways. Propranolol is highly lipophilic, in that it rapidly diffuses through the blood-brain barrier to bind directly to beta-adrenoceptors in the central nervous system, as well as in the periphery^40,41^. Conversely, as bisoprolol is highly lipophobic and therefore cannot cross the blood-brain barrier, it acts solely in the periphery^42^. If the modulations in interoceptive processes we observe in this study resulted from manipulation of the central beta-adrenergic pathway, we would expect to see significant upregulation of these processes under propranolol compared to bisoprolol, which are not evident in this study. We therefore conclude that these significant increases in cardioceptive threshold and respiroceptive precision result primarily from the modulation of peripheral beta-adrenoceptor pathways.

Our findings further suggest that beta-blockade enhances interoceptive sensitivity and metacognitive awareness by altering the autonomic balance of ascending cardiac signals toward a greater parasympathetic dominance. Both beta-blockers reduced heart rate, increased HRV, and lowered sympathovagal balance, indicating a reduced sympathetic drive and enhanced vagal control. Sympathetic activity accelerates the heart rate and reduces beat-to-beat variability, producing a more tonic and less distinct afferent signal. Vagal activity, however, slows the heart and introduces finely regulated, phasic fluctuations in cardiac intervals^43^. Beta-blockade may therefore improve the fidelity of afferent cardiac signals, increasing their signal-to-noise ratio against the background of endogenous cognitive fluctuations^44^.

Several studies have examined pharmacological manipulations of interoception, typically employing non-selective noradrenergic manipulations. Recent work has demonstrated that yohimbine, a non-selective α_2_-adrenoceptor antagonist which also activates dopaminergic and serotonergic pathways, increases cardiac sensitivity and activation in the insular cortex, a key region for interoceptive processing^45,46^. Previous findings have also suggested that noradrenaline amplifies the salience and detectability of cardiac signals, rather than its blockade. For example, Khalsa et al.^47^ found that infusion of isoproterenol, a peripheral noradrenergic agonist, increased the correlation between subjective cardiac ratings and drug-induced cardiac acceleration. These findings, together with our own, may suggest an inverted U-shaped relationship between noradrenergic stimulation and interoception, such that both agonism and antagonism of beta-adrenoreceptors increases sensitivity to visceral signals. However, these studies relied on non-selective drug manipulations and focused on exclusively cardiac measures of interoception. Our study here expands upon these findings by employing both highly selective and non-selective pharmacological agents to examine and compare specific contributions of cardiac versus central beta-adrenoceptors in more domain-general interoceptive processes spanning multiple organ systems.

In the respiratory domain, the selective enhancement of respiroceptive precision without changes in sensitivity or metacognition suggests that the consistency of respiratory judgments, rather than their detectability or evaluation, may be modulated by peripheral beta-adrenergic pathways. Similar to our cardioceptive findings, we did not find a significant difference in respiroceptive precision between propranolol and bisoprolol. Since bisoprolol is highly cardioselective, one might expect that any respiratory effect would be stronger under propranolol, which is non-selective and binds to pulmonary beta-2 adrenoceptors with similar affinity to cardiac adrenoceptors^42^. This increase in respiroceptive precision may therefore be a downstream effect of the reduced heart rate and increased signal-to-noise ratio resulting from both beta-adrenergic manipulations, in that participants’ respiratory perceptions may be subject to reduced influence from cardiac noise. Future studies may benefit from the combination of direct respiratory manipulation (e.g., using breathing exercises) with beta-blockade to better disentangle these mechanisms.

Interoception, as an inherently noisy perceptual domain, is markedly susceptible to demand characteristics, with prior studies showing a strong influence of suggestibility and subjective belief on the perception of bodily states^48,49^. Our study thus marks a significant advance in that we could here use a fully blinded design, which is further strengthened by our multiple positive control analyses which confirmed that participants exhibited no awareness of pharmacological effects. Crucially, neither drug significantly altered mood or anxiety relative to pre-drug baseline measures, as assessed by PANAS and STICSA scores. Subjective interoceptive measures, assessed via the IBQ, also remained stable under beta-blockade. These findings reinforce that the observed changes in sensitivity, precision, and metacognition reflect genuine shifts in interoceptive processing.

Our study further benefits from the use of novel interoceptive tasks that improve upon classical methods, enhancing methodological rigour and overcoming key limitations. Traditional cardiac interoception tasks^50–52^ do not account for subjective prior beliefs about heart rate or dissociate interoceptive constructs such as bias, sensitivity, and precision^53^. Our tasks build on prior approaches^54,55^ by incorporating adaptive Bayesian modelling (Psi-adaptive Bayesian psychophysical method^34,56^), enabling the efficient individualised estimation of interoceptive sensitivity, precision, and metacognitive awareness. These methods allow for more precise stimulus control and facilitate sophisticated modelling of interoceptive processes, illuminating the specific mechanisms by which noradrenaline controls bodily awareness.

Interoceptive dysfunction is a transdiagnostic feature of psychiatric illness, with disruptions in sensitivity and metacognitive awareness reported in anxiety, depression, bipolar disorder, psychosis, and substance use disorders^3,26,57^ (for reviews see ^2,58^), potentially arising from challenges in interoceptive precision specifically^17,29,59^. Previous studies have also examined the effect of various neurotransmitters, including noradrenaline and serotonin, on interoception in the context of psychiatry^60^. Beta-blockers have also been investigated more generally as potential treatments for anxiety and post-traumatic stress disorders, although their efficacy and relevant mechanisms of action have thus far been unclear^61,62^.

Our findings extend previous work by demonstrating that beta-blockade can selectively enhance interoceptive processes, improving both cardiac sensitivity and metacognition in an organ- and symptom-specific manner. The selective enhancement of respiratory precision observed here, further suggests that peripheral beta-adrenergic signalling may selectively stabilise respiratory perception. As respiratory precision has been previously linked to anxiety disorders, beta-blockers demonstrate strong therapeutic potential for mitigating aberrant breathlessness in anxiety and pulmonary disorders^63^. Similarly, major depressive disorder has been frequently associated with both reduced HRV and impaired cardiac sensitivity^64,65^, highlighting a potential role for peripheral beta-blockers as a novel intervention target. More broadly, our results support frameworks advocating for targeted interoceptive interventions^2,58^, by demonstrating that sensitivity, precision, and metacognition can be independently modulated through pharmacological strategies targeting the brain or body. Given their established safety profile and widespread clinical use, peripherally-selective beta-blockers may offer an overlooked therapeutic avenue for treating interoceptive dysfunction, particularly in psychiatric conditions where autonomic dysregulation contributes to cardiac or respiratory symptomatology.

In conclusion, our study demonstrates that peripheral beta-blockade differentially modulates interoceptive mechanisms across cardiac and respiratory domains. Through hierarchical Bayesian psychometric modelling, we reveal domain-specific roles of peripheral beta-adrenoceptors in modulating interoceptive sensitivity, precision, and awareness, refining our understanding of how noradrenaline supports interoception. Our findings therefore provide a novel perspective on how distinct mechanisms of neuromodulation may differentially impact multiple interoceptive domains, and highlight the role of peripheral beta-adrenergic pathways in the modulation of brain-body interaction on a multi-organ level.

## Methods

### Participants

The study was performed as a single-centre trial between May 2023 and December 2023 at the Centre for Functionally Integrative Neuroscience (CFIN), Aarhus University. From 60 healthy young adult volunteers initially recruited, 57 were assigned for randomisation, 57 received at least one intervention and 52 completed all components of the study. Data from one participant was removed from all analyses due to an adverse drug reaction, and data from one participant was removed due to poor task compliance. Therefore 50 participants (mean age: 25.2 ± 5.1 S.D., 29 females) were included in our analyses. For the final analyses of interoceptive tasks, 49 participants were included for the HRDT, and 48 were included for the RRST (see **Supplementary Figure S4** for CONSORT flowchart). Four participants had incomplete data for the HRDT for one of their study visits; data for their remaining visits were included in our analyses.

Recruitment occurred via public advertisements at Aarhus University and Aarhus University Hospital, social media, and via an institutional online database. All participants provided written informed consent according to the declaration of Helsinki (2013) and received remuneration for their participation. This project was approved by the local Ethics Committee (De Videnskabsetiske Komitéer For Region Midtjylland, ID: 102977). Right-handed participants aged between 20 and 40 years were recruited. Only participants with no history of cardiological, neurological or psychiatric diagnoses, or contraindications to any study interventions were included in the study (detailed inclusion and exclusion criteria in **Supplementary Methods S4**). Participants also underwent a brief cardiac examination by a clinician prior to participation to ensure cardiac health.

### Pharmacological Intervention

The study followed a randomised, placebo-controlled, counter-balanced, double-blind design. Each participant attended three separate study visits scheduled at least five days apart to allow for complete drug washout. At each visit, a single oral dose of either 40 mg propranolol, 10 mg bisoprolol, or matched placebo (inert sugar pills) was administered 140 minutes prior to the interoception tasks. The time point was selected to optimise peak plasma concentration of both propranolol and bisoprolol during the tasks^66,67^.

The selected doses were informed by established clinical guidelines and prior research. Both 40 mg propranolol and 10 mg bisoprolol represent the minimal effective starting doses commonly prescribed for hypertension in standard clinical practice (as recommended by www.indlaegssedler.dk), ensuring their safety, tolerability, and suitability for human research. Propranolol was further chosen based on its extensive use in prior studies investigating its effects on cognition and brain activity at this dose^8,9,68–70^. In contrast, bisoprolol has not previously been studied in the context of cognition or brain function, offering a novel opportunity to explore its influence on interoceptive and cognitive processes.

All medications were oral tablets matched in appearance and delivery method to ensure double-blinding of the entire study team. Balanced, block-wise randomisation, and blinding were performed by an experienced researcher not involved in data collection. Confirmation of participant blinding efficacy is described in **Supplementary Methods S5**.

### Analysis of Electrocardiograph Data from Resting State Magnetoencephalography

To obtain metrics of HRV for each participant, we analysed ECG data recorded while participants rested with their eyes open during an MEG scan, which took place approx. 65 minutes before participants completed the interoceptive tasks. The corresponding resting-state MEG results will be reported elsewhere. R peak detection (using the ECGLAB fast R peak detection algorithm) and quality control were conducted using the HEPLAB toolbox^71^, and HRV metrics were calculated using the HRVAS toolbox^72^, both in MATLAB. All ECG data were visually inspected for artefacts and erroneous R peak detection. All data included in this analysis contained at least two minutes of continuous cardiac recording. Pairwise Student’s *t* tests were then conducted using JASP to examine differences between mean heart rate (BPM), RMSSD, and LF/HF ratio between each of the three drug groups. Five sessions (one session each for five different participants: two placebo, two bisoprolol, and one propranolol session) could not be included in this analysis due to missing data.

### Psychological and Mental Health Measures

To characterise subclinical mental health symptoms, we assessed ADHD, depression, and anxiety symptoms using the Adult ADHD Self-Report Scale (ASRS), the Beck Depression Inventory-II (BDI-II), and the Beck Anxiety Inventory (BAI), respectively. These assessments were conducted at the beginning of the study prior to any interventions through online surveys. Additionally, participant demographics such as height, weight, age, sex, and BMI, were recorded (**Table 1**).

**Table 1:** Participant Demographics.

| Variable | Mean (± SD) |
| --- | --- |
| Age (Mean ± SD) | 25.2 ± 5.1 |
| Height (cm) | 173.5 ± 10.3 |
| Weight (kg) | 68.6 ± 13 |
| BMI | 22.7 ± 2.8 |
| ADHD (ASRS) | 2.5 ± 1.4 |
| Anxiety (BAI) | 10.1 ± 6.4 |
| Depression (BDI II) | 6.7 ± 6.4 |
| Sex | 29 (58%) Female, 21 (42%) Male |

*Note: BMI = Body Mass Index. BAI = Beck Anxiety Inventory, BDI-II = Beck Depression Inventory II, ASRS = Adult ADHD Self-Report Scale.*

Drug effects on mental state variables were assessed with self-report scales at baseline (immediately prior to drug administration) and 70 minutes post-drug administration. These included the 11 somatic items (items 1, 2, 6, 7, 8 12, 14, 15, 18, 20, 21) contained in the State-Trait Inventory for Cognitive and Somatic Anxiety (STICSA), and the Positive and Negative Affect Scale (PANAS). We also examined interoceptive self-beliefs associated with cardiac and respiratory domains, using the Interoceptive Belief Questionnaire (IBQ) (**Supplementary Table 1**).

### Interoceptive Psychophysics Test Battery

To quantify interoception in the cardiac and respiratory domains, we employed a battery of validated psychophysical measures, namely the HRDT^32^ and the RRST^33^ (**Figure 1**). These tasks used adaptive Bayesian methods to estimate key psychophysical parameters, including interoceptive sensitivity (threshold) and precision (slope). Participants completed 60 trials in each task. Interoceptive task order was counterbalanced between participants but kept consistent for each participant across all study visits.

Both tasks featured a two-interval forced-choice design and were carefully designed to minimise non-interoceptive cues (e.g., auditory or visual feedback) that might influence performance. Custom-designed setups ensured robust and reproducible delivery of sensory stimuli, with inter-trial intervals and task durations optimised to maintain participant engagement and minimise fatigue. The adaptive Bayesian approaches used in both tasks provide efficient and reliable estimates of key psychophysical parameters while reducing the number of trials needed compared to traditional staircase methods. Hierarchical ordered beta regression was also applied to confidence and response accuracy data from both tasks to examine how beta-blockade may influence metacognition (see *Statistical Analyses)*.

#### Heart Rate Discrimination Task

In the HRDT, participants were asked to silently attend to their own heart rate without physically checking their pulse during a 5 s ‘heart listening’ phase (**Figure 1B**). Participants then heard a series of tones at varying tempos, presented at a fixed frequency above or below the current heart rate, and were asked to indicate whether these feedback tones were faster or slower than their own heart rate using the left and right computer mouse buttons. Participants then provided a confidence rating assessing their judgement accuracy using a visual analogue scale (VAS) ranging from ‘Guess’ to ‘Certain’. For the duration of the task, the participant’s heart rate was monitored continuously by a Nonin soft-clip pulse oximeter placed on the fourth finger of the left (non-dominant) hand. The auditory feedback tones were dynamically adjusted using the Psi adaptive algorithm^34^, which efficiently estimated the threshold (sensitivity) and slope (precision) of the psychometric function in units of delta-beats per minute (Δ-BPM). The HRDT was administered through the PsychoPy package in Python^73^.

### Respiratory Resistance Sensitivity Task

The RRST is a psychophysical task in which participants were asked to take two successive inhalations through a computer-controlled respiratory circuit (**Figure 1C**). A resistive load was randomly applied to one of these breaths (the stimulus) via a precision step motor compressing the circuit tubing, whereas the other breath was without resistive load. The magnitude of the stimulus on each trial was also controlled using the Psi adaptive method, estimating threshold, slope, and lapse rate parameters. After taking the two breaths, participants were asked to indicate whether they found the first or second breath to be more difficult using the left and right computer mouse buttons. Participants then provided a confidence rating assessing their judgement accuracy using a VAS ranging from ‘Guess’ to ‘Certain’. To eliminate auditory cues from the motorised apparatus, participants wore noise-cancelling headphones playing continuous white noise. The Psi-adaptive algorithm was used to adjust the resistance levels across trials to estimate the psychometric threshold (sensitivity to resistance) and slope (precision). The RRST was administered via Psychtoolbox and the Palamedes toolboxes in MATLAB^56,74^.

## Statistical Analyses

### Effect of Drugs on Mean and Standard Deviation of BPM during the HRDT

As a positive control analysis, we investigated whether the mean BPM and standard deviation (SD) BPM acquired during the HRDT differed between drug groups, using pairwise Student’s *t* tests.

### Effect of Drugs on Affect, Anxiety, and Subjective Interoceptive Beliefs

As a further positive control analysis, we examined whether beta-blockers were associated with any changes in the self-reported affect (PANAS) and somatic anxiety (STICSA), using repeated measures ANOVAs, with the following within-subject factors: drug (placebo, propranolol, bisoprolol), and time (baseline, 70 min post-drug). We also examined how beta-adrenergic blockade may be associated with participants’ subjective interoceptive beliefs before and after completing the interoceptive tasks, using a cumulative link mixed model for ordinal regression with the following within-subject factors: drug (placebo, propranolol, and bisoprolol) and time (before versus after completion of interoceptive tasks). Pairwise comparisons were assessed using Tukey’s HSD post-hoc test.

### Hierarchical Bayesian Modelling of Interoceptive Psychophysics

We applied a hierarchical Bayesian model to simultaneously estimate the psychophysical function at the group and single subject level, thus allowing for a parsimonious model of both population and individual level estimates^75,76^.

For the HRDT, we modelled the binary decision (i.e., “is the tone faster or slower than your heart rate?”) as a function of the difference between the frequency of auditory tones and the heart rate (beats per minute, Δ*_BPM_*) of the participant using a cumulative normal distribution with three subject-specific parameters: threshold (interoceptive sensitivity, α), slope (interoceptive precision, β) and lapse rate (λ). The cardioceptive psychophysical model is as follows:

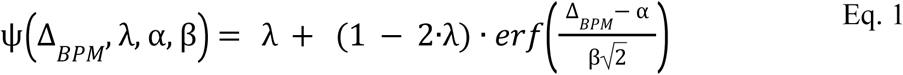

For the RRST we applied the same model but instead employing a cumulative Weibull function, and modelled the accuracy of the binary decision (“was the first or second breath more difficult?”) as a function of the difference between the resistive loads for the two breaths, estimating the same three subject-specific parameters as with the HRDT. The respiroceptive psychophysical model is as follows, where *Res* denotes the resistive load:

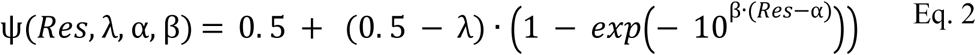

To account for pharmacological manipulation, we allowed the three model parameters to vary across manipulations by establishing the placebo condition as the reference level and each of the pharmacological conditions to induce a difference from this reference. Additionally, we controlled for the differential effect of participants’ average resting heart rate in the model for the HRDT (see **Supplementary Methods S2-3** for detailed methods).

To obtain samples from the joint posterior distribution we used Stan, a No-U-Turn Hamiltonian Monte Carlo Markov Chain Monte Carlo sampler^77^. Each model was initialised with 4 chains, 2500 warm-up samples, 2500 samples, adapt-delta of 0.99 and a max tree depth of 12. Each model was investigated for convergence by ensuring no divergences, satisfactory chain mixing, Rhat values of < 1.05^78^, and at least 400 effective samples for group level parameters. For each model we selected a set of priors that contained a feasible range of behavioural responses and that were also in accordance with previously published results of these tasks^32,33^. Crucially we selected priors for all difference parameters between placebo and both drug interventions to be unbiased (i.e., normal distributions with mean 0). We assessed statistical significance by determining whether the 95% highest density interval of the difference parameters crossed zero (i.e., placebo > bisoprolol difference, for example). One participant was rejected from our analyses of the HRDT, and two participants were rejected from our RRST analyses, based on visual inspection of the raw data. For a full set of priors on all group level parameters see **Supplementary Tables 2–3**.

### Ordered Beta Regression on Confidence Ratings

To investigate the effects of beta-adrenergic antagonism on metacognitive awareness during cardiac and respiratory interoception, we employed ordered beta regression^79^. This method is well-suited for modelling confidence ratings, which are continuous but bounded between the extremes. The ordered beta distribution works as a continuous mixture distribution with four main distributional parameters, μ, ϕ, *k*1 and *k*2.

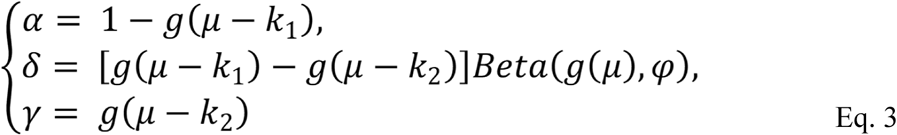

k1 and k2 serve as cut points that determine the number of extreme values, i.e., 0 and 1 respectively. ϕ is the precision parameter of beta distribution parameterised with a mean (µ) and precision parameters. Here, we parameterise the mean entailing a parsimonious shift in the whole distribution, such that higher means also entail higher proportions of ‘1’ responses.

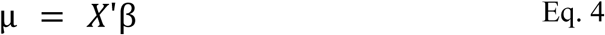

Where X is the matrix of predictor variables and β a vector of parameters. The ordered beta regression framework allowed us to estimate both metacognitive bias (average confidence per drug condition) and metacognitive awareness (sensitivity of confidence to accuracy), while controlling for the subject-specific random effects.

For the HRDT, confidence ratings were modelled as a function of drug condition (placebo, propranolol, bisoprolol), task accuracy (correct/incorrect), and the z-scored listening heart rate on each trial. For the RRST, confidence ratings were modelled as a function of drug condition, task accuracy, and stimulus intensity (i.e., % respiratory resistance). The random effects structure for both tasks included random intercepts and slopes for stimulus intensity (heart rate for the HRDT and % resistance for the RRST) and response accuracy. Post-hoc pairwise comparisons were conducted using the emmeans package in R to interrogate the significant drug-by-accuracy interactions. Tukey’s method was applied to control for multiple comparisons.

## Acknowledgements

This research is financially supported by a Lundbeckfonden Fellowship (R272-2017-4345) and a European Research Council Grant (ERC-2020-StG-948788) awarded to MGA. FF is supported by a European Research Council Grant (ERC-2020-StG-948838) and a Lundbeckfonden Experiment Grant (R436-2023-991). The funding sources were not involved in the study design, collection, analysis, interpretation, or writing of the manuscript. TUH is supported by a Sir Henry Dale Fellowship (211155/Z/18/Z; 211155/Z/18/B; 224051/Z/21) from Wellcome & The Royal Society and a grant from the Jacobs Foundation (2017-1261-04). We also thank Daniel Kluger for his valuable inputs in the editing of this manuscript.

## Disclosures

All authors declare no conflicts of interest.

## Supplementary Material

### S1: No meaningful effect of beta-blockade on subjective interoceptive beliefs

As an additional positive control, we examined the effects of beta-blockade on subjective interoceptive beliefs, using the Interoceptive Belief Questionnaire (IBQ, see **Supplementary Table 2**). The IBQ assesses aspects of affect and sensitivity in the domains of heartbeat and breathing, and was administered both immediately before and after participants completed the interoceptive tasks. For heartbeat sensitivity, breathing sensitivity, and breathing affect, there were no significant differences between drug conditions either before or after the interoceptive task battery (all contrasts: *p* > .1; ordinal regression model). There was a significant difference in heartbeat affect under bisoprolol (but not propranolol) compared with placebo (*p* = .04), in that heartbeat affect was significantly lower under bisoprolol prior to the interoceptive tasks compared with placebo, but not post-task. However, this effect did not survive post-hoc pairwise comparison (*p* = .0996; Tukey’s HSD). This therefore suggests that neither propranolol nor bisoprolol exerted a meaningful effect on interoceptive beliefs, and that interoceptive beliefs were not significantly influenced during completion of the interoceptive tasks.

### S2: Reduced mean heart rate under beta-blockade during HRDT

We also assessed the effects of propranolol, bisoprolol, and placebo on mean heart rate recorded during the HRDT using linear mixed effects modelling. As expected, mean heart rate was significantly reduced under propranolol and bisoprolol compared to placebo (propranolol: *t*(47) = 10.5, *p* < .001; bisoprolol: *t*(47) = 10.4, *p* < .001). There was no significant difference in mean heart rate between propranolol and bisoprolol (*t*(46) = 1.8, *p* = .17). These findings confirm the effect of both propranolol and bisoprolol on reducing overall heart rate during the resting state ECG is also consistent during the HRDT.

### S3: Control Analysis

To bolster our findings, we performed a control analysis to account for the heart rate of each participant recorded during the HRDT in our hierarchical parameter estimation. All results obtained for the HRDT include this control, and our control pipeline is described below.

We conceptualised the control analysis in the following manner. Each participant attended one placebo session and two sessions with pharmacological manipulation. These pharmacological interventions may have altered their mean heart rate (HR) by some degree on an individual level (heart rate difference pre- versus post-intervention; ΔHR). However, such interventions may also alter the threshold of the psychometric function (Latent Threshold Difference; ΔLT) for the HRDT. Here, we simulated the underlying thresholds (alphas) and set the HR for the simulated agents to 55 bpm. We then obtained their trial-by-trial stimulus alpha values, to enable the hierarchical model fitting of all simulated agents, and estimated the difference in alpha (i.e., the difference between the agents’ HR during the task and their latent thresholds), both with and without controlling for HR.

We simulated data with varying parameter values, using values of 0-10 for ΔLT and 0-15 for ΔHR, running for 100 iterations per parameter value set. We used this to further examine the difference in threshold between real data and both sets of simulated data; i.e., real data compared with controlled and uncontrolled simulations.

#### Data simulations with control analyses generate parameter estimates closer to real data compared with uncontrolled simulations

We then simulated HRDT data both with and without this control analysis, and found that the controlled simulated data was significantly closer to the real data compared with the simulated data in the absence of control analysis. By repeating this process using multiple values for ΔHR and ΔLT, we found that this remains the case across multiple parameter values, and the divergence of the uncontrolled simulated data increases as values for ΔHR increase. Values for ΔLT did not significantly affect the difference between the real data and both sets of simulated data, controlled and uncontrolled.

### S4: Full Participant Inclusion and Exclusion Criteria

#### Inclusion Criteria

- Age 20-40 years
- Participant is able and willing to give informed consent
- Normal or corrected-to-normal vision
- In case of incidental findings, the participant consents that we may share their data with a specialist who may contact the participant if deemed medically necessary
- Good English language skills
- Must consent to and pass multi-panel drug tests at each visit (urine tests)
- Body Mass Index (BMI) in the healthy-to-normal range (18.5 - 30)

#### Exclusion Criteria

- Regular use of medications, with the exception of over-the-counter antihistamines or
- contraceptives
- History of neurological diseases of the motor and sensory system that may interfere with the
- conduct of the study
- History of psychiatric diseases, disorders or conditions
- History of cardiac diseases, conditions or disorders
- Family history of heart failure
- History of any substance abuse disorder
- History of chronic pain, diabetes, cancer, or diseases of the liver, kidney, intestinal or
- cardiovascular systems
- Contraindications to propranolol or bisoprolol (listed below)
- Pregnancy, possible pregnancy, planned pregnancy, or lactation
- Jet lag or sleep deprivation
- MRI safety conditions not met (i.e., participant was not claustrophobic, and participant had no metals in/on the body that could not be removed for scanning)
- Any other reason to exclude the participant according to judgement by the investigator

### S5: Participant Blinding Efficacy

To examine the extent of potential unblinding, we evaluated whether participants were able to correctly guess which drug they had been administered. At the end of each session, participants were asked to guess which drug they thought they had been given. The accuracy of these guesses was then compared against the actual drug administered to the participant during that session. We performed a group-level binomial test to determine whether the overall accuracy of guesses across participants was significantly different from chance (⅓).

At the group level, the binomial test revealed that participants were not able to guess the drug with accuracy significantly greater than chance (95% CI [0.27; 0.43], *p* = .74, observed correct guessing rate = 35%).

### Figures and Tables

**Supplementary Figure S1:**
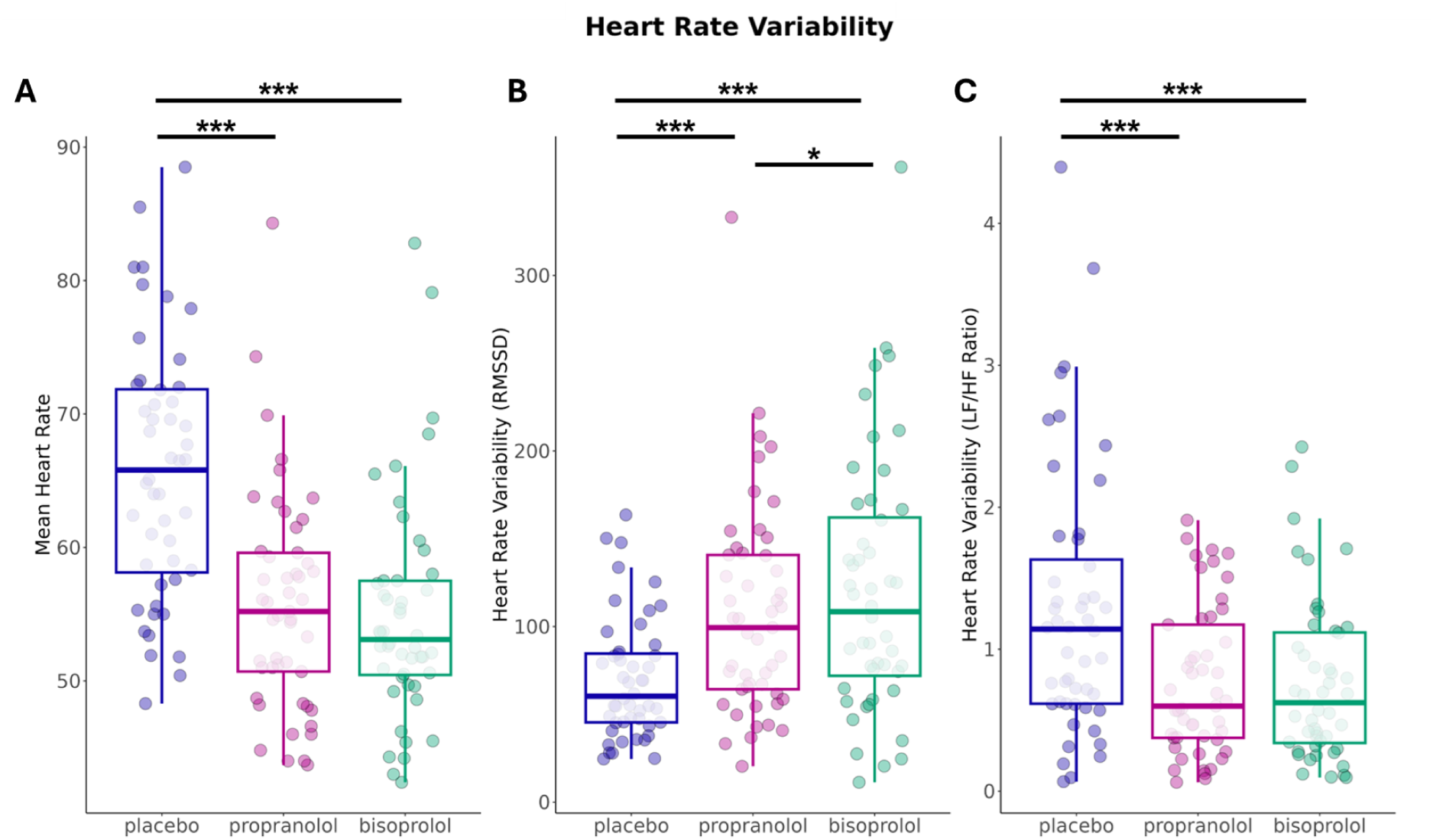
Effects of drug condition on mean heart rate and HRV metrics (RMSSD and LF/HF ratio). Mean heart rate (**A**), and HRV (RMSSD (**B**) and LF/HF ratio (**C**) calculated from resting state ECG data. *** = *p* < .001, * = *p* < .05. Error bars = range excluding outliers. Both propranolol and bisoprolol significantly reduced mean heart rate and LF/HF ratio compared to placebo (\*\*\**p* < .001), with no significant differences between propranolol and bisoprolol. RMSSD was significantly increased under propranolol and bisoprolol compared with placebo (\*\*\**p* < .001). RMSSD was also significantly increased under bisoprolol compared with propranolol (\**p* = .029).

**Supplementary Figure S2:**
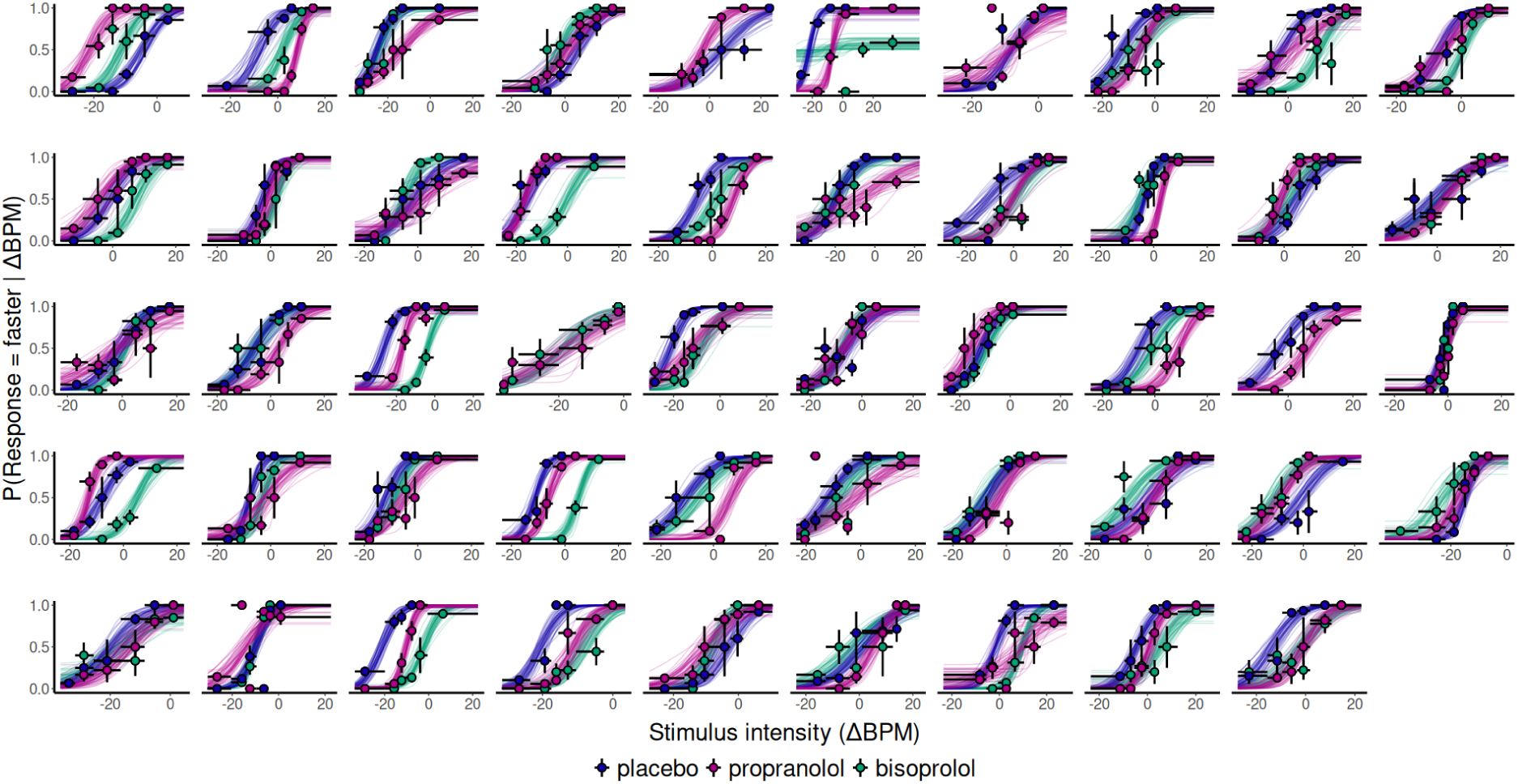
50 draws of individual predictive participant-level psychometric functions for each drug condition in the HRDT. Each facet represents a different participant with the colours representing each drug condition. Points display binned raw responses; seven equally sized bins were created for each subject. Error bars on the points display the standard error of the bin in the vertical direction and the size of the bin in the horizontal direction.

**Supplementary Figure S3:**
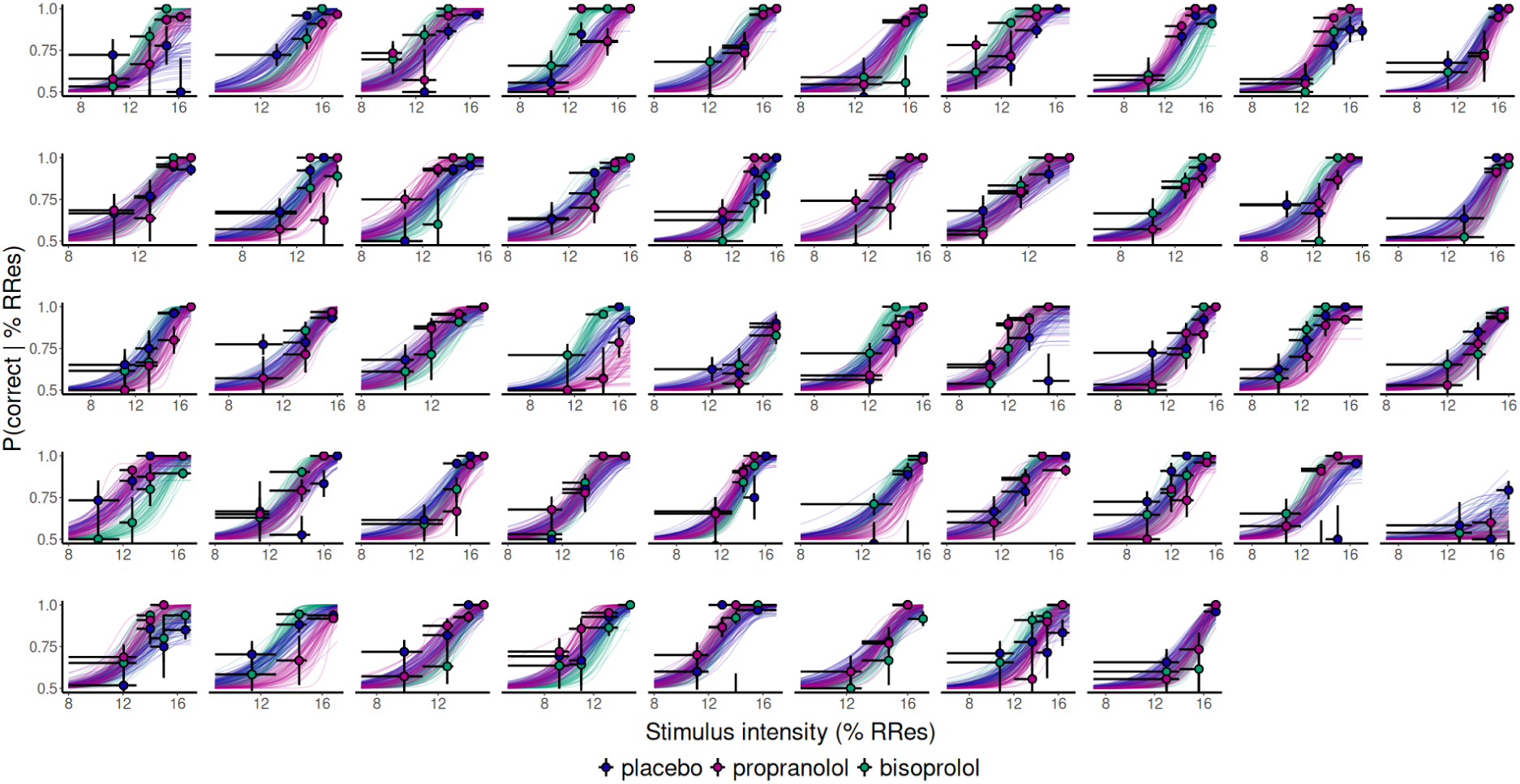
50 draws of individual predictive participant-level psychometric functions for each drug condition in the RRST. Each facet represents a different participant with the colours representing each drug condition. Points display binned raw responses; five equally sized bins were created for each subject. Error bars on the points display the standard error of the bin in the vertical direction and the size of the bin in the horizontal direction.

**Supplementary Figure S4:**
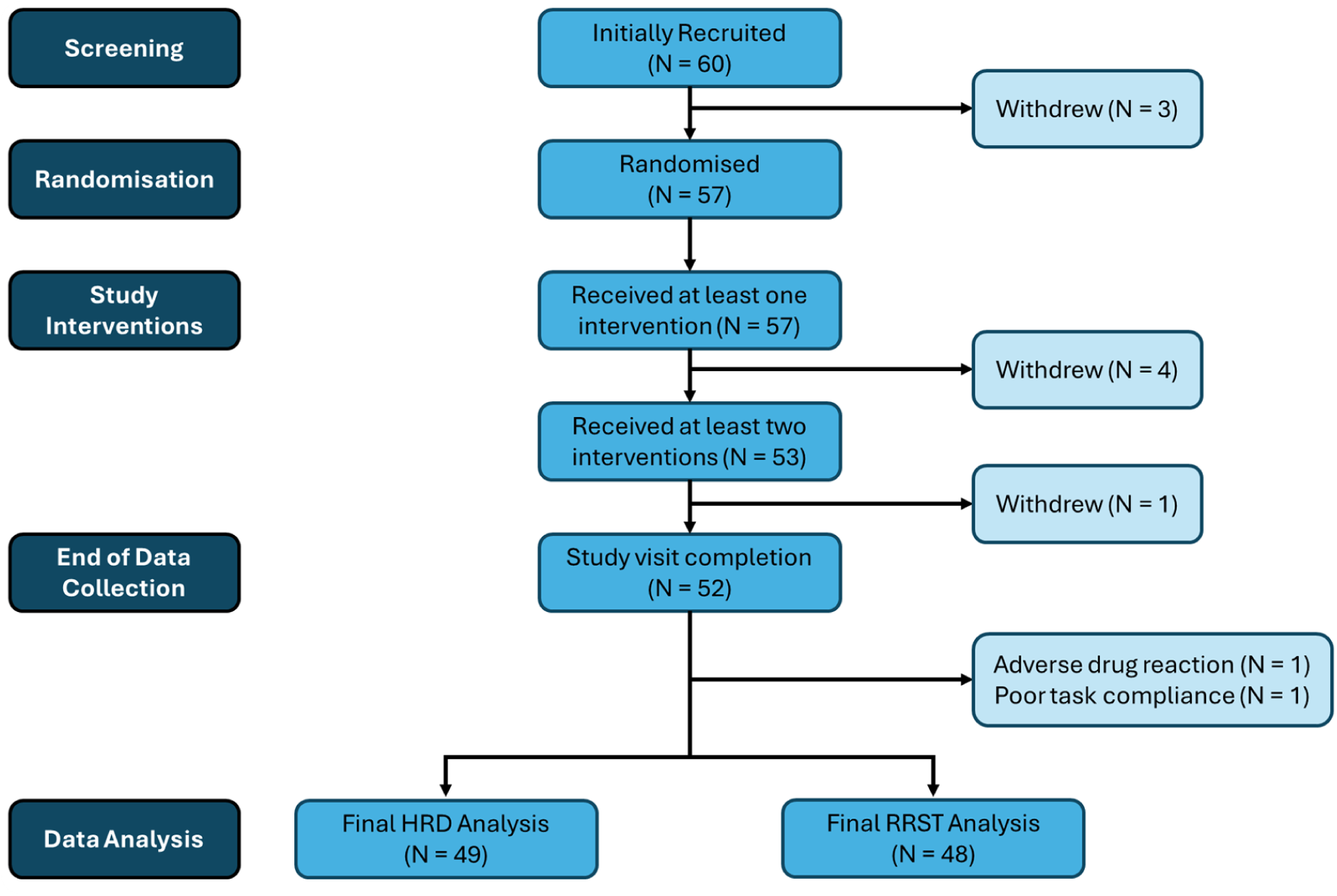
CONSORT flow chart displaying the progress of all participants throughout the study. Of the 60 initially recruited volunteers, 49 participants were included in the HRDT data analysis, and 48 participants were included in the RRST analysis.

### Supplementary tables

**Supplementary Table 1:** Interoceptive Belief Questionnaire: Modalities and Response Scales.

| **Modality** | **Item Text** | **Response Scale** |
| --- | --- | --- |
| Breathing Sensitivity | I am generally sensitive to the sensations of my own breathing. | 1 (Strongly Disagree) - 10 (Strongly Agree) |
| Heartbeat Sensitivity | I can generally tell how fast my heart is beating. | 1 (Strongly Disagree) - 10 (Strongly Agree) |
| Heartbeat Affect | When you focus on the sensation of your heartbeat, how does it make you feel? | 1 (Extremely Disturbed) - 7 (Extremely Relaxed) |
| Breathing Affect | When you focus on the sensation of your breathing, how does it make you feel? | 1 (Extremely Disturbed) - 7 (Extremely Relaxed) |

*Note:* Items 1 and 2 are belief-based questions rated on a 1 to 10 scale. Items 3 and 4 assess affective responses to interoceptive focus, rated on a 1 to 7 scale.

**Supplementary Table 2:**
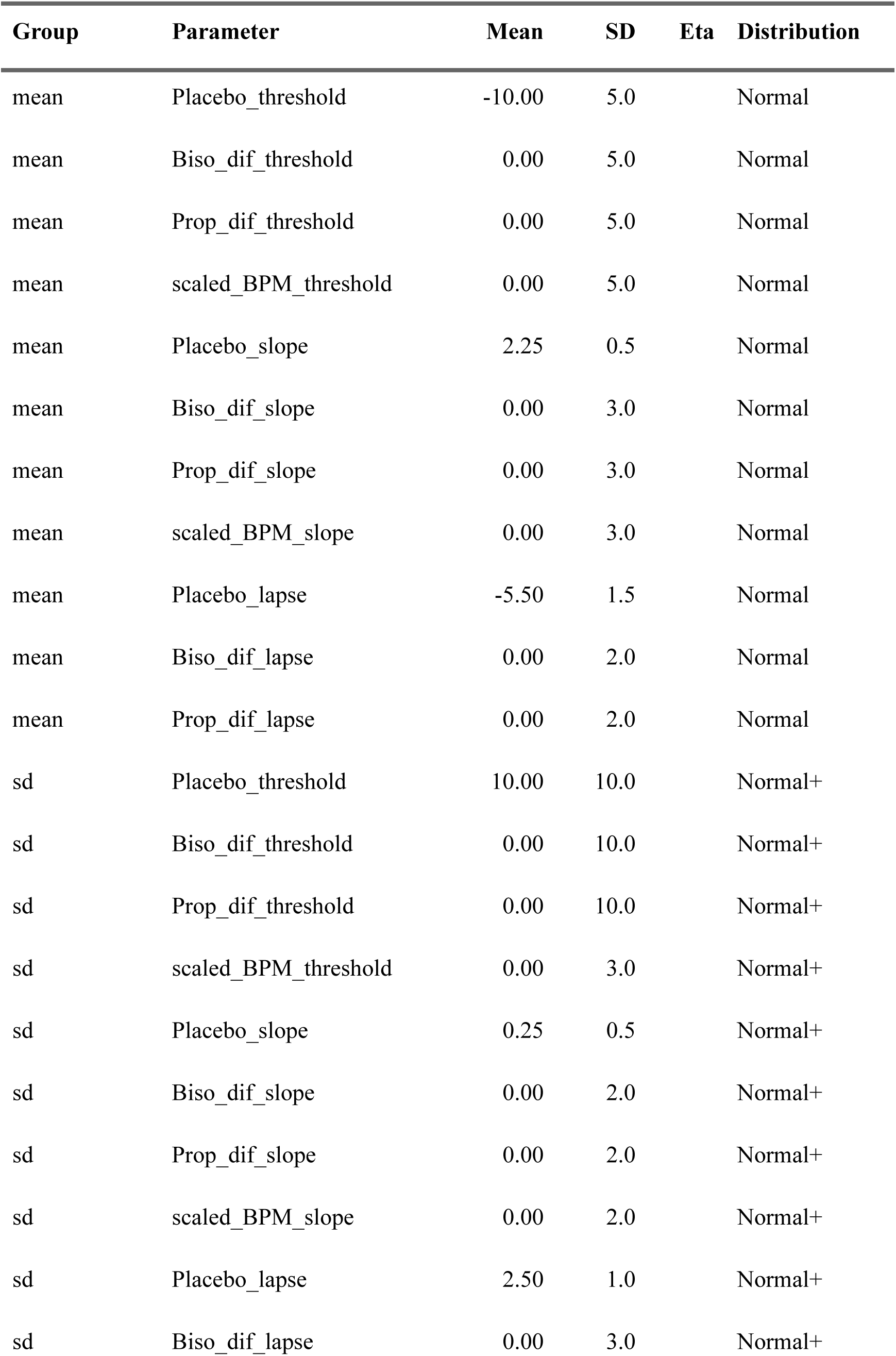
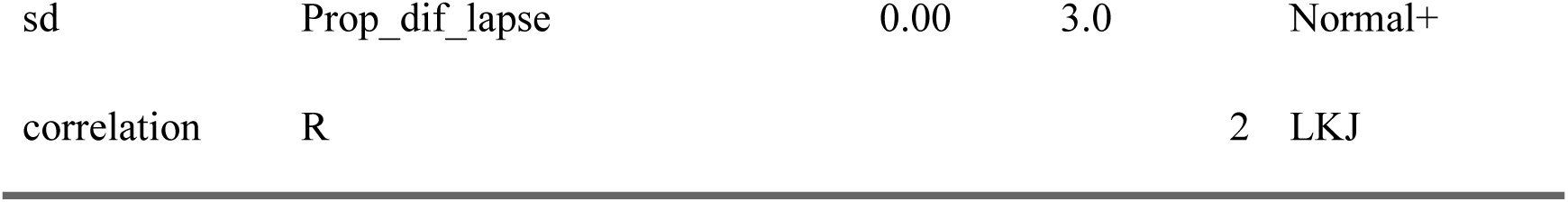
Full set of priors for the HRDT.

**Supplementary Table 3:** Full set of priors for the RRST.

| Group | Parameter | Mean | SD | Eta | Distribution |
| --- | --- | --- | --- | --- | --- |
| mean | Placebo_threshold | 10 | 10.0 |  | Normal |
| mean | Biso_dif_threshold | 0 | 5.0 |  | Normal |
| mean | Prop_dif_threshold | 0 | 5.0 |  | Normal |
| mean | Placebo_slope | 0 | 3.0 |  | Normal |
| mean | Biso_dif_slope | 0 | 3.0 |  | Normal |
| mean | Prop_dif_slope | 0 | 3.0 |  | Normal |
| mean | Placebo_lapse | -4 | 3.0 |  | Normal |
| mean | Biso_dif_lapse | 0 | 1.5 |  | Normal |
| mean | Prop_dif_lapse | 0 | 2.0 |  | Normal |
| sd | Placebo_threshold | 0 | 5.0 |  | Normal+ |
| sd | Biso_dif_threshold | 0 | 5.0 |  | Normal+ |
| sd | Prop_dif_threshold | 0 | 5.0 |  | Normal+ |
| sd | Placebo_slope | 0 | 3.0 |  | Normal+ |
| sd | Biso_dif_slope | 0 | 3.0 |  | Normal+ |
| sd | Prop_dif_slope | 0 | 3.0 |  | Normal+ |
| sd | Placebo_lapse | 0 | 3.0 |  | Normal+ |
| sd | Biso_dif_lapse | 0 | 3.0 |  | Normal+ |
| sd | Prop_dif_lapse | 0 | 3.0 |  | Normal+ |
| correlation | R |  |  | 2 | LKJ |

## Notes

### Competing Interest Statement

The authors have declared no competing interest.

